# Nitrogen starvation leads to TOR kinase-mediated downregulation of fatty acid synthesis in the algae *Chlorella sorokiniana* and *Chlamydomonas reinhardtii*

**DOI:** 10.1101/2023.07.31.551344

**Authors:** Jithesh Vijayan, Sophie Alvarez, Michael J Naldrett, Amanda Maliva, Nishikant Wase, Wayne R. Riekhof

## Abstract

The accumulation of triacylglycerol (TAG) as a storage compound in eukaryotic algae has been the subject of extensive studies over the last 50 years. The model industrial alga *Chlorella sorokiniana* accumulates TAG and other storage compounds under nitrogen (N)-limited growth. Previously we used transcriptomics to explore the regulation of TAG synthesis in *C. sorokiniana*. Surprisingly, our analysis showed that the expression of several key genes encoding enzymes involved in plastidic fatty acid synthesis are significantly repressed. Metabolic labeling with radiolabeled acetate showed that *de novo* fatty acid synthesis is indeed downregulated under N-limitation. Likewise, inhibition of the Target of Rapamycin kinase (TOR), a key regulator of metabolism and growth, decreased fatty acid synthesis. We compared the changes in proteins and phosphoprotein abundance using a proteomics and phosphoproteomics approach in *C. sorokiniana* cells under N-limitation or TOR inhibition and found extensive overlap between the N-limited and TOR-inhibited conditions. We also identified changes in the phosphorylation levels of TOR complex proteins, TOR-kinase and RAPTOR, under N-limitation, indicating that TOR signaling is altered. Our results indicate that under N-limitation there is significant metabolic remodeling, including fatty acid synthesis, mediated by TOR signaling. We find that TOR-mediated metabolic remodeling of fatty acid synthesis under N-limitation is conserved in the chlorophyte algae *Chlorella sorokiniana* and *Chlamydomonas reinhardtii*.

## INTRODUCTION

Algae as a feedstock for biofuels and other bioproducts has been investigated over the last 50 years, owing to the potential for high biomass yield, accumulation of large quantities of neutral lipids, sequestration of carbon dioxide, and capability of growth on marginal lands and in brackish wastewater. (Hu et al., 2008). Triacylglycerol (TAG) derived from biological sources can be transesterified and blended with crude oil-derived diesel for burning in Diesel engines, and can serve as a feedstock to produce jet fuel and other hydrocarbon-based products (Allen et al., 2018; Beal et al., 2021). Microalgae accumulate TAG under certain stress conditions, such as nutrient limitation (Zhu et al., 2016) or chemical stressors (Wase et al., 2017), and the mechanism by which macronutrient limitation induces TAG accumulation has been the subject of many studies for the past five decades (Bollmeier and Sprague, 1989; Wase et al., 2018; Li-Beisson et al., 2019). Fatty acids for TAG synthesis can be derived either from *de novo* synthesis or by membrane remodeling, i.e., the process of channeling fatty acids and glycerol from other membrane glycerolipids, such as monogalactosyldiacylglycerol and phosphatidylcholine (Li et al., 2012; Yoon et al., 2012; Li-Beisson et al., 2019; Chen and Wang, 2021; Vijayan et al., 2021; Young and Shachar-Hill, 2021).

In the plant lineage, fatty acid synthesis occurs in the chloroplast, and Acetyl CoA Carboxylase (ACCase) and Fatty Acid Synthase complex (FAS) enzymes are similar to the prokaryotic multiple-subunit “type II” synthases (Post-Beittenmiller et al., 1992). Carboxylation of acetyl-CoA to malonyl-CoA catalyzed by ACCase is the first committed step in fatty acid biosynthesis. FAS complex uses malonyl-CoA to sequentially add 2-carbon units to the nascent fatty acid, ultimately forming a 16-18 carbon chain (Wakil et al., 1983). Fatty acid synthesis is a major sink of ATP and reducing equivalents and is hence highly regulated (Salie et al., 2016).

Target of Rapamycin (TOR) is a phosphatidylinositol-3-kinase like serine/threonine kinase that is present in all eukaryotes and is known to regulate metabolism in response to a variety of cues, such as hormones, nutrients, and cellular or organismal energy status. It is known to be a nexus connecting nutrient sensing, growth, and metabolism, acting in the regulatory module of SnRK1/TOR (Robaglia et al., 2012). In yeast and humans, there are two TOR complexes, TORC1 and TORC2. Both complexes have TOR-kinase and LST8 as common members, and the presence of RAPTOR (TORC1) or RICTOR (TORC2) determines the identity and function of the complex (Laplante and Sabatini, 2012). In plants and algae TORC2 is absent as a plant homolog of RICTOR has not been identified (Pérez-Pérez et al., 2017).

*Chlorella sorokiniana* is a microalga belonging to the family Trebouxiophyceae. It is an industrial microalga with high growth potential (Lizzul et al., 2018) with the ability to grow on municipal and agricultural wastewater (Bohutskyi et al., 2015). *C. sorokiniana* genome is sequenced and well characterized and annotated (Arriola et al., 2018). In this article, we present evidence for regulation of fatty acid synthesis by TOR signaling under N-limitation conditions in the microalgae *C. sorokiniana* and *Chlamydomonas reinharditii*.

## RESULTS

### Nitrogen starvation in C. sorokiniana leads to a decrease in the rate of fatty acid biosynthesis

In a previous transcriptomic analysis of *Chlorella sorokiniana* cultures under N-limitation, we observed that many genes encoding key enzymes in various steps of fatty acid synthesis are downregulated as depicted in Fig. S1A-B (Vijayan et al., 2021). Transcript abundances of Ketoacyl-ACP-Synthase 1 (KAS1), 3 (KAS3), Hydroxyacyl-ACP-Dehydratase (HAD1) and Enoyl Reductase (ENR1) are significantly decreased. These enzymes are components of the plastidic FAS complex as outlined in the introduction. The observed downregulation of these genes prompted us to hypothesize that fatty acid synthesis is decreased in N-limited cultures. To test this hypothesis, we carried out metabolic labeling studies with ^14^C-acetate under N-replete and -limited conditions after 24 hours of growth. The experiment was carried out with 10^7^ cells for each condition and experimental replicates. Under N-limitation, incorporation of ^14^C-acetate into fatty acid is significantly decreased, as shown in Fig. 1A and 1B. We also observed no significant increase in total fatty acid content in cultures under N-limitation, on a per cell basis (Fig. 1C). While there is no significant increase in total fatty acid content, we observed changes in fatty acid compositions, with oleic (18:1) and linoleic acid (18:2) increasing in abundance, and linolenic acid (18:3) and its 16-carbon counterpart (16:3^Δ7,10,13^) decreasing in abundance (Fig. 1D). Incorporation of acetate into fatty acid is decreased, but the flux of existing fatty acids into TAG is increased, as shown in Fig. 1E. We found that expression of genes encoding enzymes involved in the TCA cycle are also downregulated (Fig. S1C). Expression of malate dehydrogenase, fumarase and citrate synthase 1 are significantly downregulated. This suggests that large-scale remodeling of central carbon metabolism is a component of the response to N-limitation in *C. sorokiniana*.

**Fig 1.**
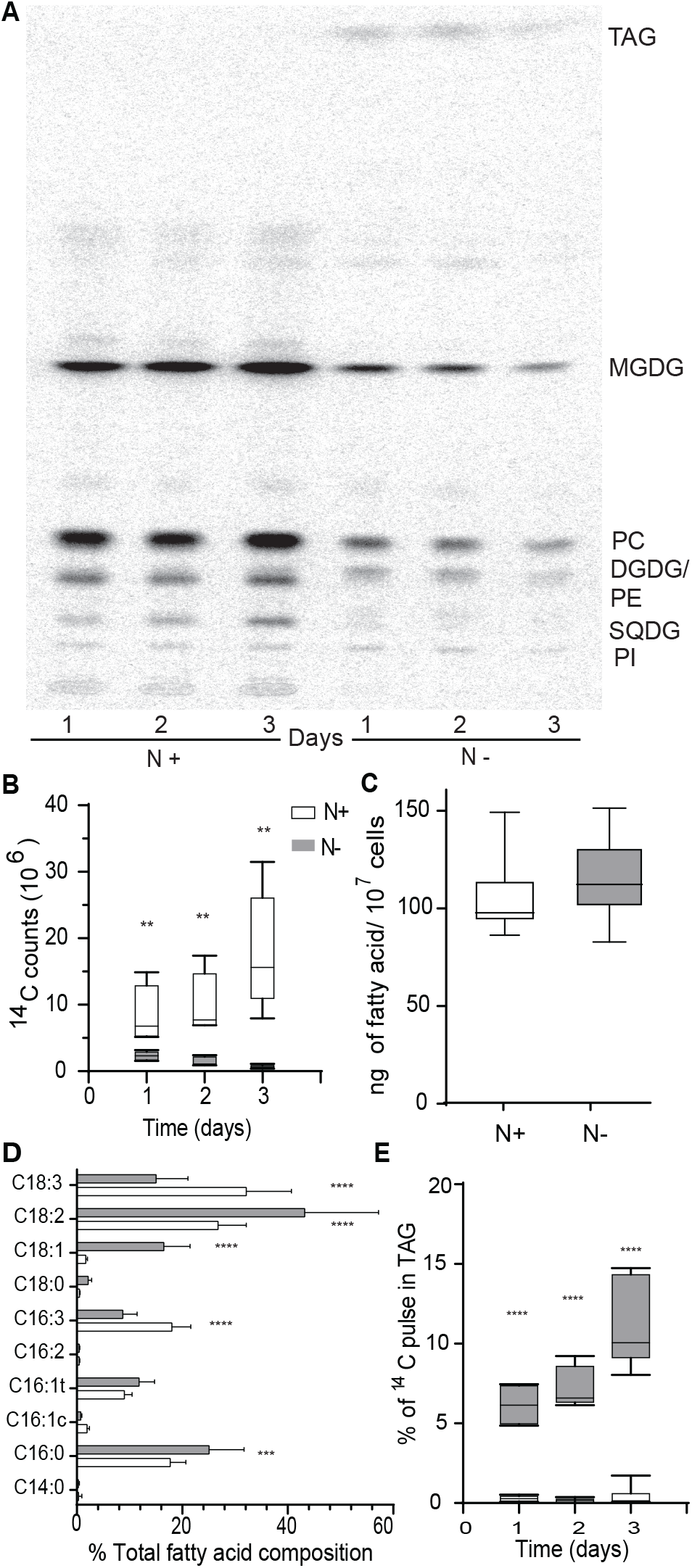
Downregulation of fatty acid under nitrogen starvation in *C. sorokiniana*. A) TLC-autoradiograph of 10^⁷^ *C. sorokiniana* cells grown in N replete (N+) and deplete (N-) media fed with ^14^C acetate after 1, 2, and 3 days of inoculation. B) Quantitative measurement of ^14^C acetate incorporated into fatty acids under nitrogen replete (N+) and deplete (N-). C)) Total Fatty Acid content of cells under nitrogen replete and deplete conditions as normalized by number of cells. D) Percent fatty acid composition of cells under nitrogen replete and deplete conditions. E) Percentage of ^14^C acetate incorporated into TAG under nitrogen replete. * indicate the statistical significance of student t test. n=9. Error bars indicate SD

### Inhibition of TOR kinase leads to decreased fatty acid biosynthesis, similar to N-limitation

The relationship between TOR signaling and N-metabolism is well established in the budding yeasts *Saccharomyces cerevisiae* and the fission yeast *Schizosaccharomyces pombe*. Deletion of TOR2 in fission yeast leads to N-limitation like responses (Matsuo et al., 2007). The same is true of seed plants, including *Arabidopsis thaliana*, in which N-limitation has been shown to regulate TOR activity (Liu et al., 2021). The transcriptome of *C. sorokiniana* under N-limitation also showed that the expression of RAPTOR1, a TOR complex component, is doubled (Fig. 2A). This observation and prior studies in other organisms prompted us to hypothesize that the activity of the TOR complex could be altered under N-limitation in *C. sorokiniana*.

**Fig 2.**
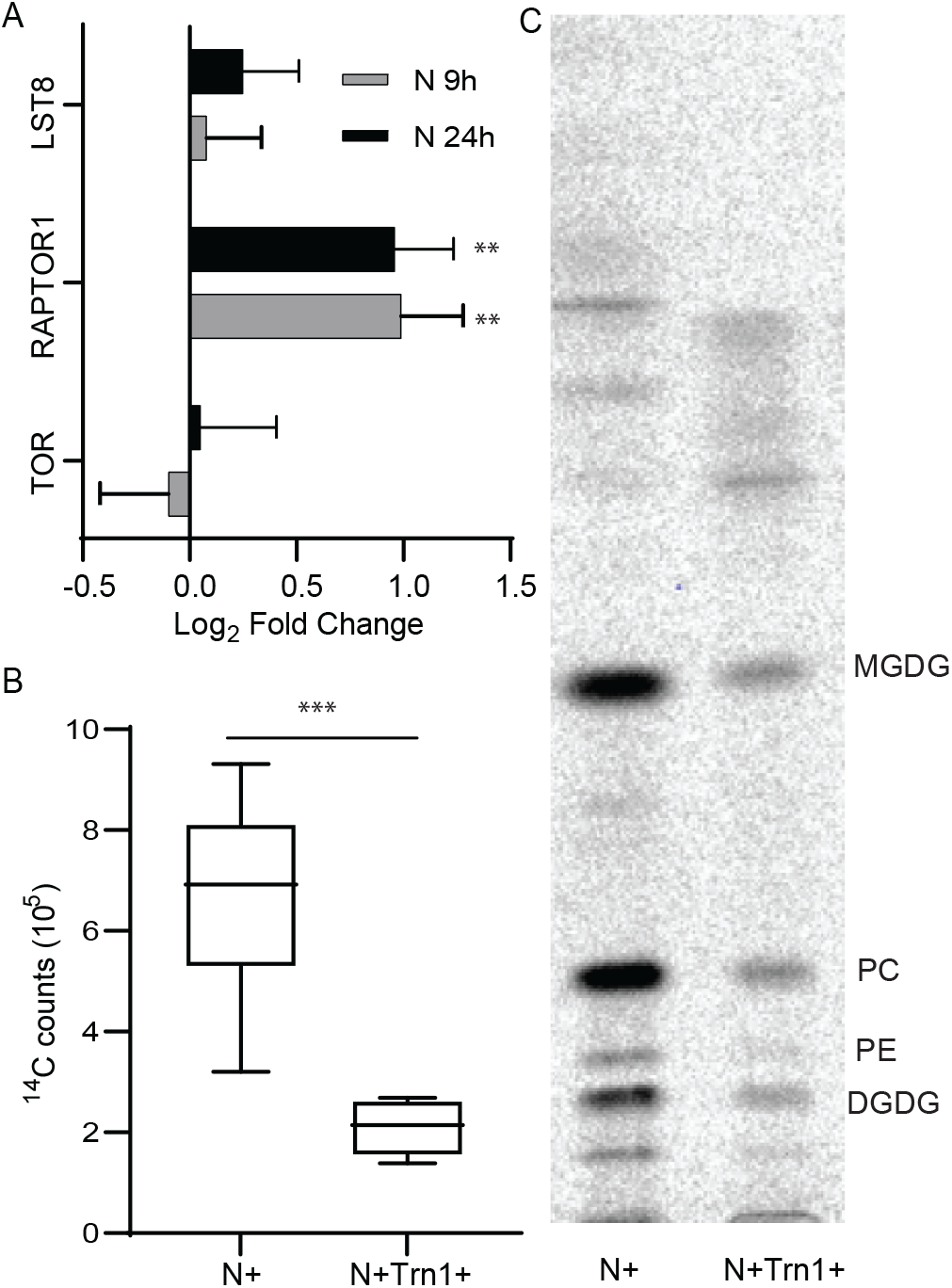
Control of fatty acid synthesis under nitrogen limitation by TOR. A) Expression levels of different known TOR complex members under nitrogen limitation conditions. B) TLC-autoradiograph of *C. sorokiniana* cells fed with ^14^C-acetate to track de novo fatty acid synthesis grown in N replete media with (N+Trn1+) or without (N+) TOR inhibitor-torin1 C). Quantitative measurement of ^14^C-acetate incorporation into fatty acid as a measure of de novo fatty acid synthesis when fed with (N+Trn1+) or without (N+) Torin1. * indicate the statistical significance of student t test. n=9. Error bars indicate SD

TOR signaling is known to affect lipogenesis via the SREBP1 pathway in humans (Porstmann et al., 2008). In *Saccharomyces cerevisiae*, TORC1 regulates lipid droplet formation through transcription factors such as Gln3p, Gat1p, Rtg1p, and Rtg3p (Madeira et al., 2015). Furthermore, Imamura et al. have shown that inhibition of TOR induces TAG accumulation in the microalga *Cyanidioschyzon merolae* (Imamura et al., 2016). This indicates that TOR signaling affects lipid metabolism in a number of organisms that are evolutionarily divergent, so we asked if downregulation of fatty acid synthesis under N-limitation is mediated by altered TOR signaling in *C. sorokiniana*. When *C. sorokiniana* cells grown in N-replete media were treated with rapamycin, we did not observe a robust or reproducible inhibition of growth (Fig. S2). Li et al have observed that rapamycin does not inhibit the growth of *C. sorokiniana* (Li et al., 2022). We decided to use Torin 1, a known competitive inhibitor of ATP binding of TOR kinase (Thoreen et al., 2009). Unlike Li et al., we observed that Torin 1 inhibits growth of *C. sorokiniana*. This shows that TOR signaling is likely essential for growth of *C. sorokiniana*.

When cells were fed ^14^C-acetate after 4 hours of treatment with 5 µM Torin 1 in N-replete media, we see that acetate incorporation into fatty acid is significantly decreased in comparison to the control cultures (Fig. 2B, 2C). Control cultures were cells grown in N-replete media without Torin 1. Decreased fatty acid synthesis when TOR kinase activity is inhibited by Torin 1 indicates that TOR controls fatty acid synthesis in *C. sorokiniana*.

### N-limitation alters TOR signaling

To further understand the mechanism of TOR mediated regulation of fatty acid metabolism, and relationship between TOR signaling and N-limitation, we performed a quantitative proteomic and phosphoproteomic analysis of cultures grown under these conditions. *C. sorokiniana* cultures were grown in N-replete and -limited conditions. Twenty-two hours after inoculation, an aliquot of N-replete culture was diluted to 10^7^ cells/ml and Torin 1 was added to a final concentration of 5 µM. Cultures with Torin 1 were incubated for 90 minutes before pelleting for sample preparation. N-limited and -replete cultures were harvested 24 hours after inoculation. N-replete samples were used as control for both Torin 1 treated and N-limited samples.

A total of 3424 proteins were identified of which 3237 were identified with at least ≥ 2 peptides and quantified across the three treatments. These proteins have a molecular weight ranging from 4-634 kDa. Changes in abundance were calculated, with a fold-change of 0.5 and lower considered downregulated, and 1.5 and above as upregulated, with a p-value < 0.05. The abundance of 299 and 133 proteins decreased in N-limited and Torin 1 treated samples, respectively, while 52 proteins decreased in abundance in both conditions as shown in Fig. 3A. This accounts for 17% of the total proteins downregulated in N-limitation and 39% for Torin 1 treatment. Hypergeometric probability analysis shows a significant overlap with a representation factor of 4.5 (p < 7.144e-23). Similarly, the abundance of 214 and 246 proteins increased in N-limited cells and Torin 1 treated samples respectively (Fig. 3A) and 94 proteins in both conditions which amounts to 44% of proteins in the N-limited condition. (Representation factor: 6.1; p < 1.305e-56)

**Fig 3.**
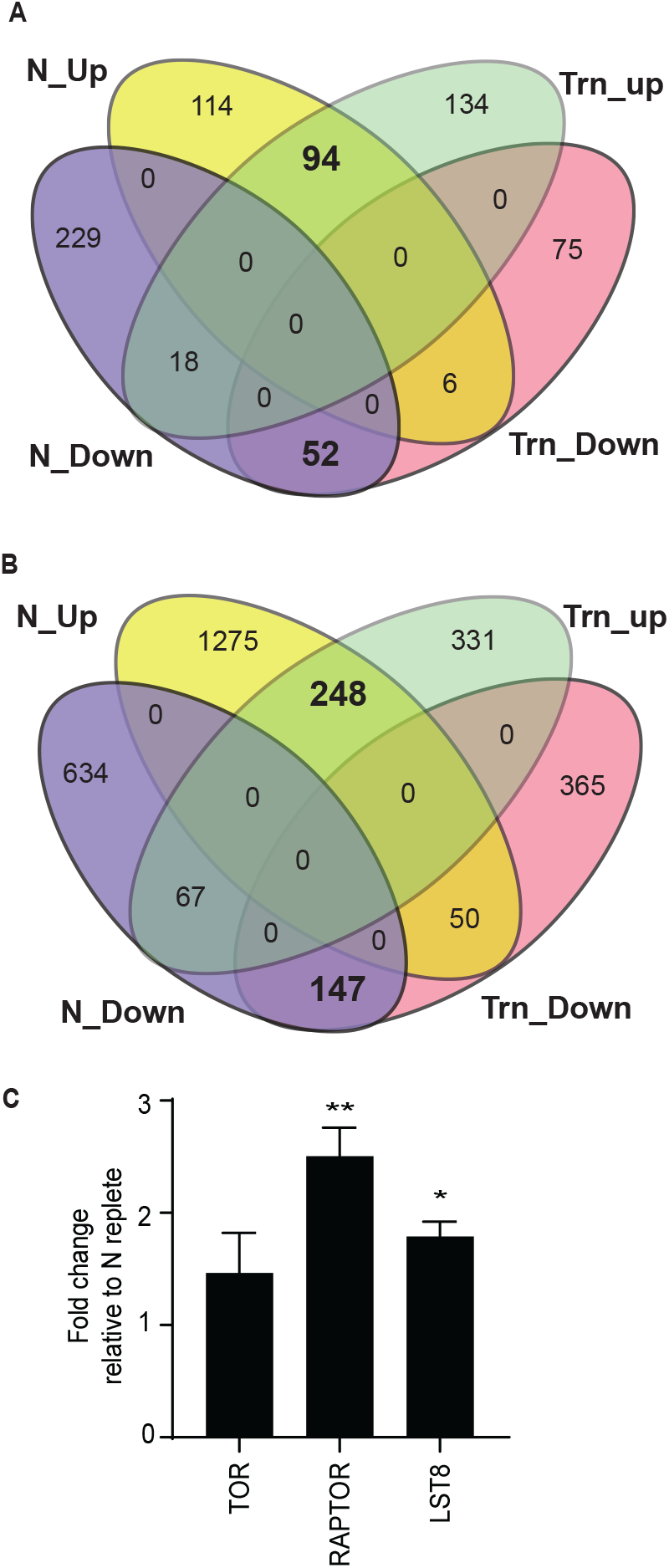
TOR signaling is affected under nitrogen limitation. A) Venn diagram showing the number of proteins that are differentially regulated under nitrogen limitation (N) and torin1 treatment (trn). Fold change is relative to control (N replete) cutoff, ≤ 0.5and ≥ 1.5. p-value ≤ 0.05 or less. (n=3). B) Venn diagram showing the number of phosphopetides that are differentially regulated under nitrogen limitation and torin1 treatment. Fold change relative to control (N replete) cutoff, ≤ 0.75and ≥ 1.5. p-value ≤ 0.05 or less. (n=3)

A total of 6477 unique phosphopeptides belonging to 1985 proteins were detected in the phosphoproteomic analysis. Fig. 3B indicates the number of peptides whose phosphorylation status was significantly (p < 0.05) decreased under N-limitation and Torin 1 treatment, with fold change cutoffs of ≥1.5 and ≤ 0.5 for N-limited samples and ≥1.5 and ≤ 0.75 Torin 1 treatment. The less stringent cutoff for the Torin 1 treatment comparison was chosen to account for the shorter incubation period of 90 minutes, versus the 24-hour N-limitation treatment. 848 phosphopeptides belonging to 549 proteins had decreased phosphorylation levels under N-limitation, while Torin 1 treatment affected 562 phosphopeptides belonging to 402 proteins. 147 phosphopeptides were commonly dephosphorylated under both conditions which is 17% of peptides in N-limitation samples whose phosphorylation is decreased (Representation factor: 2.0; p < 1.632e-18). 1573 peptides had increased phosphorylation under N-limitation and 248 of these peptides were common between Torin 1 treated cells and N-limited cells (Fig. 3B) (Representation factor: 1.6; p < 2.636e-17). There is a substantial and statistically significant overlap of proteome and phosphoproteome between N-limitation and TOR inhibition, indicating that TOR activity is affected under N-limitation.

Protein levels of known TOR complex members, RAPTOR and LST8 are increased in nitrogen starved cells (Fig. 3C). We also observed differential phosphorylation of RAPTOR and TOR kinase (Table 1). In TOR kinase, T1195 and S1581 were hyperphosphorylated while S1209 and S2439 were hypo phosphorylated. S2439 is in a motif pSppr. This motif is adjacent to the catalytic domain and conserved in many different algae (Fig. 4). This serine is found to be phosphorylated in *C. reinharditii* as well (Wang et al., 2014). Further, we observed that S504, S510, S514, S521 and S523 of RAPTOR were hypophosphorylated under N limitation. These residues are in TOR-interacting FAT domain. The effect of these phosphorylations on the kinase activity and protein-protein interactions are beyond the scope of this work, however we hypothesize that these modifications affect the functioning of the TOR complex. Conservation of the pSppr motif near the C-terminus of the TOR-kinase in different algal lineage raises the possibility that TOR activity could be controlled by differential phosphorylation of this residue by a conserved kinase whose identity is yet unknown.

**Fig 4.**
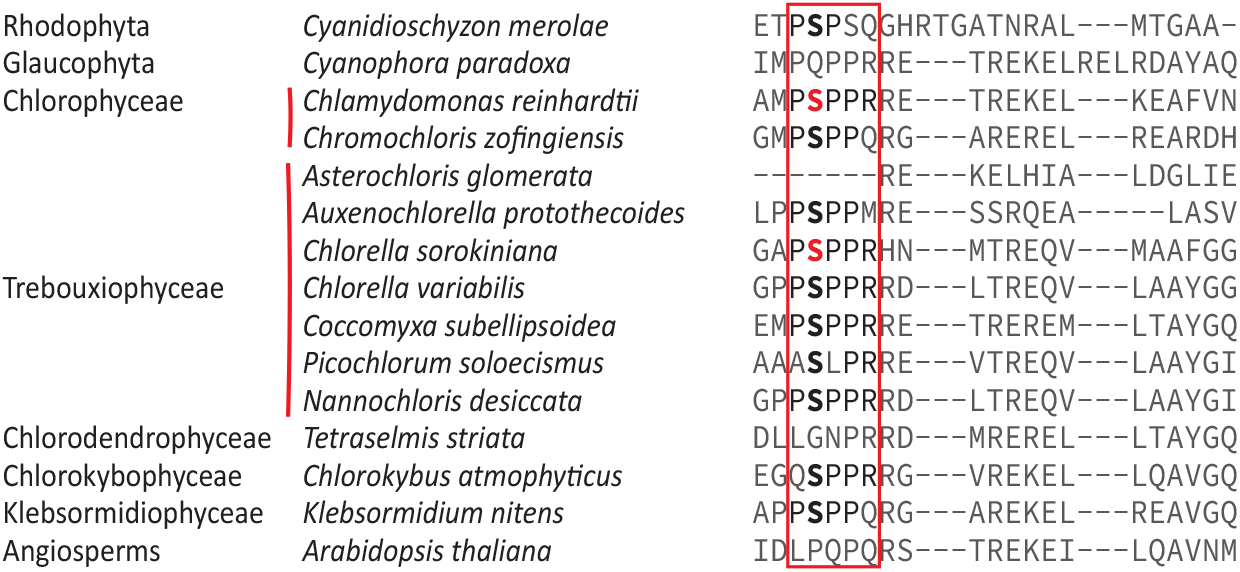
Conservation of pSppr motif in TOR kinase of different microalgae. Multiple sequence alignment of TOR kinase protein from different microalgae highlighting (red box) the conserved pSppr motif in the C-terminus. Serine in red indicates known phosphorylation of the residue.

**Table 1:**
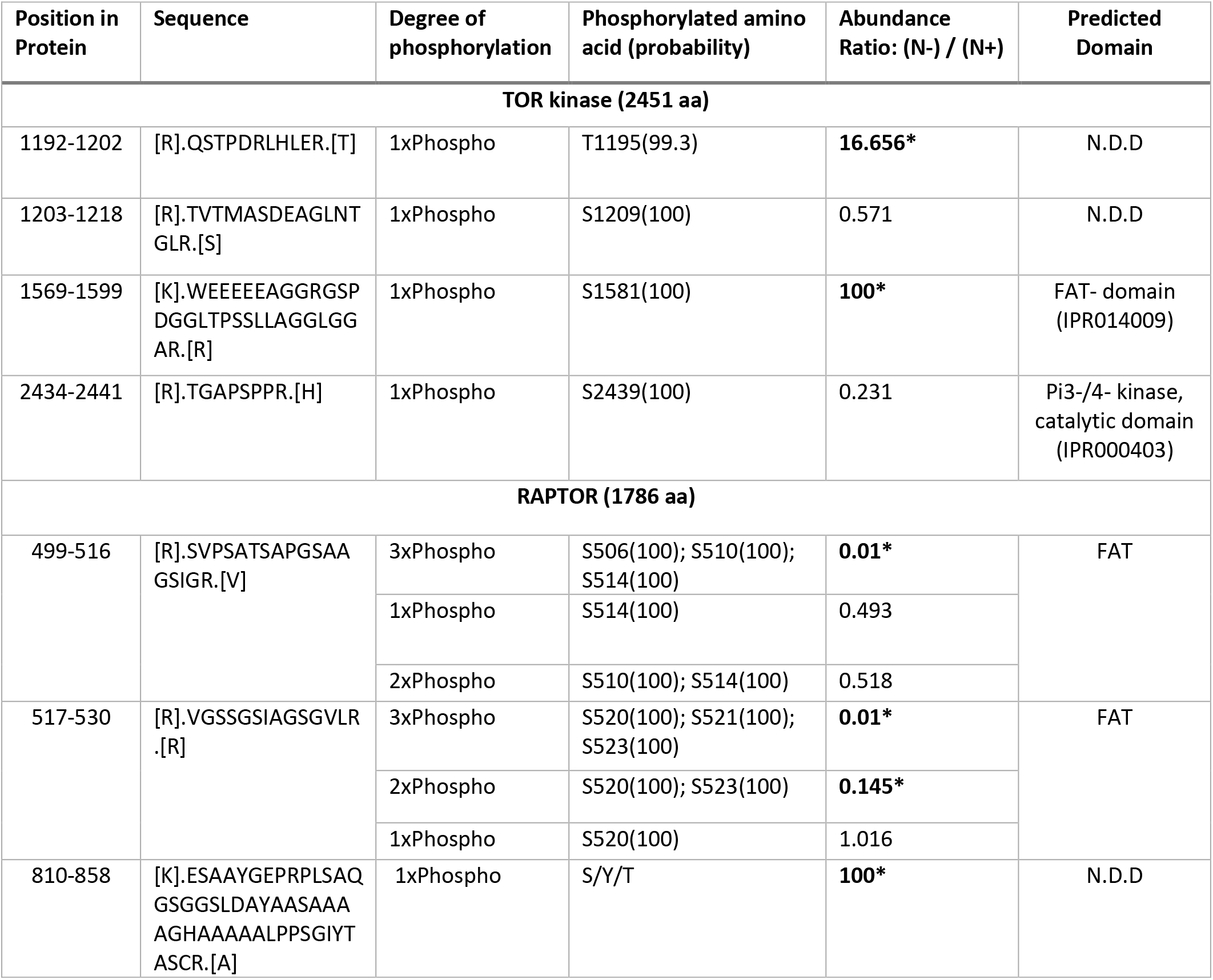
Phosphorylation status of TOR complex proteins. Ratios of 0.01 and 100 are arbitrary fold changes to indicate that the peptides are not detected in the treated conditions and in control samples, respectively. N.D.D: No Domain Detected. * indicates statistical significance.

Robust and statistically significant similarity between the proteome and phosphoproteome under N-limitation and Torin 1 treatment, and changes in the phosphorylation status of TOR complex components in the N-limited condition suggest that TOR signaling is altered under N-limitation.

### Effect of N-limitation and TOR inhibition on ACCase and FAS complex components

We observed that protein levels of the carboxyltransferase component of the plastidic ACCase complex is decreased under both N-limitation and TOR inhibition (Fig. 5A). 3-oxoacyl-ACP reductase of the FAS complex is also significantly downregulated under N-limitation (Fig. 5B). We hypothesize that these changes in the protein levels affect the rate of fatty acid synthesis. It should be noted that the expression of many genes encoding FAS complex members are downregulated under N-limitation, but the protein levels do not follow the pattern. We did not observe any phosphorylation of plastidial-ACCase or FAS complex members. Nevertheless, reduction in protein levels of a core component of the ACCase and of 3-oxoacyl-ACP reductase (FAS) would be expected to decrease the rate of fatty acid synthesis. Additionally, in a transcriptomic study of *C. sorokiniana* cells treated with the TOR kinase inhibitor AZD8055, Li *et al*. observed that the expression of all FAS complex members are downregulated (Li et al., 2022). This indicates that TOR controls fatty acid synthesis at both the transcriptional level, and in regulating protein abundance via changes in translation or protein turnover.

**Fig 5.**
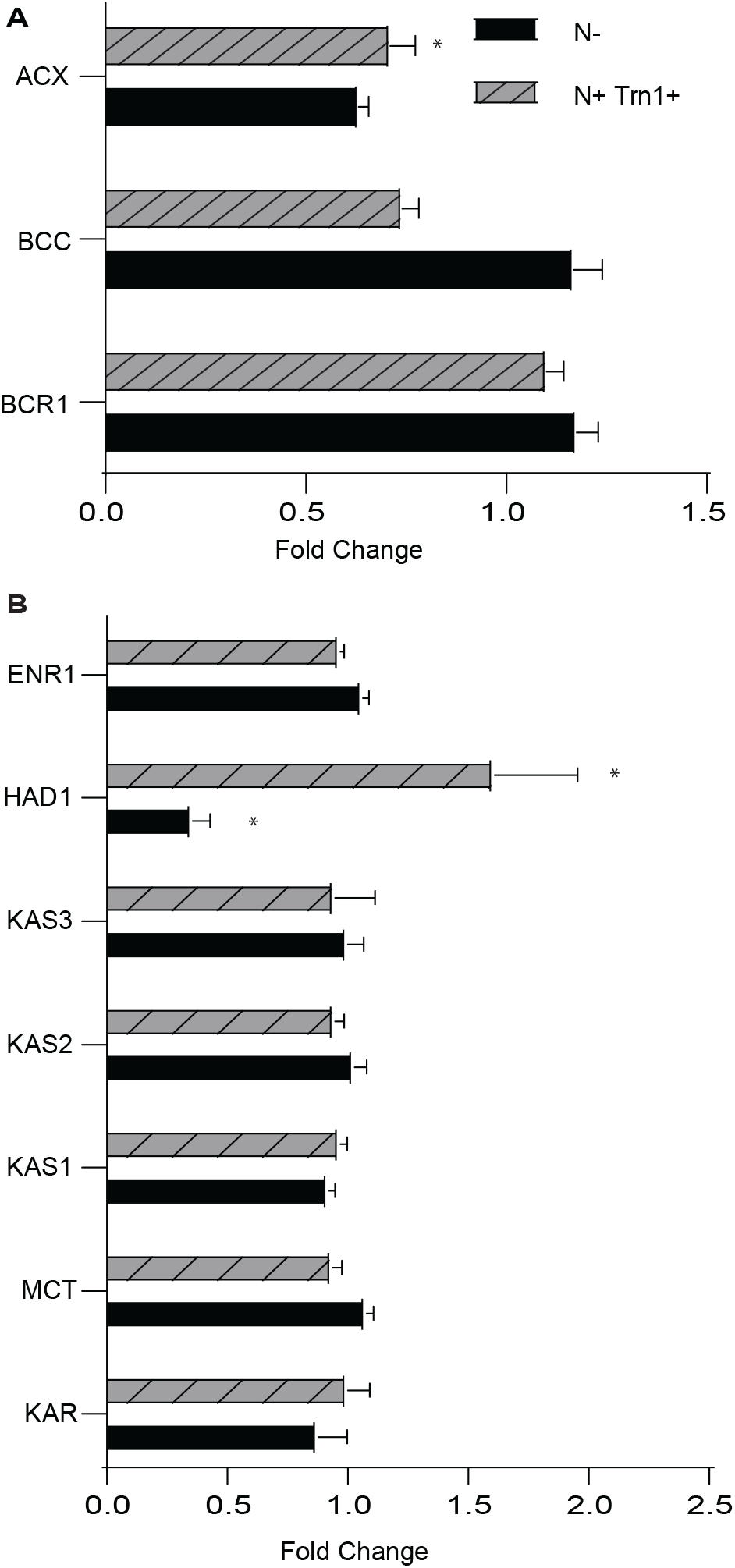
Protein levels of ACCase and FAS complex members. A) Protein levels of different components of ACCase complex members under nitrogen limitation (N-) and Torin 1 treatment (N+Trn1+). BCR1: Biotin carboxylase, BCC: Acetyl-CoA biotin carboxyl carrier protein, ACX: Carboxyltransferase. B) Protein levels of different enzymes involved in the process of fatty acid synthesis. KAR: 3-Oxoacy-ACP reductase, MCT: Malonyl-CoA: ACP transacylase, KAS: 3-Ketoacyl-ACP-synthase, HAD: 3-Hydroxyacyl-ACP dehydratase, ENR: Enoyl-[acyl carrier protein] reductase. Error bars indicate SD (n=3).

### TOR mediated regulation of fatty acid synthesis under N-limitation in Chlamydomonas reinhardtii

*C. sorokiniana* and the well-studied model microalgae *C. reinhardtii* are members of the taxa Chlorophyta. Therefore, we enquired if similar mechanisms of TOR-signaling mediate control of fatty acid synthesis under N-limitation in *C. reinhardtii.* For this, we first queried the published transcriptome of *C. reinhardtii* under N-limitation (Schmollinger et al., 2014) and found that expression of all the genes encoding components of the plastidic ACCase and FAS complexes were downregulated as early as 4 hours after inoculation in nitrogen limited media (Fig. S3A, B). This indicates that there is indeed a transcriptional mechanism for downregulation of fatty acid synthesis under N-limitation. Next, we took advantage of the RNAseq data published by Kleessen *et al*. to investigate if inhibition of TOR activity by rapamycin affects the transcript levels of these genes (Kleessen et al., 2015). While the expression of genes encoding BCC1, BCC2, KAS2, and KAS3 decrease to a very small extent, we see that BCR1 and KAR1 expression decreases by half after 4 hours of rapamycin treatment (Fig. S3C, D). Thus, TOR inhibition affects fatty acid synthesis at a transcriptional level in *C. reinhardtii*, similar to *C. sorokiniana*. To query if the transcriptional downregulation of these genes indeed affect fatty acid synthesis in *C. reinharditii* under N-limitation or TOR inhibition, we carried out a ^14^C-acetate labeling experiment. We used either 5 µM Torin 1 or 1 µM rapamycin to inhibit TOR activity. Incorporation of ^14^C-acetate was significantly decreased under N-limitation and Torin 1 treatment (Fig. 6A, B), thus indicating that TOR-mediated control of fatty acid synthesis under N-limitation is conserved between *C. sorokiniana* and *C. reinhardtii*.

**fig 6.**
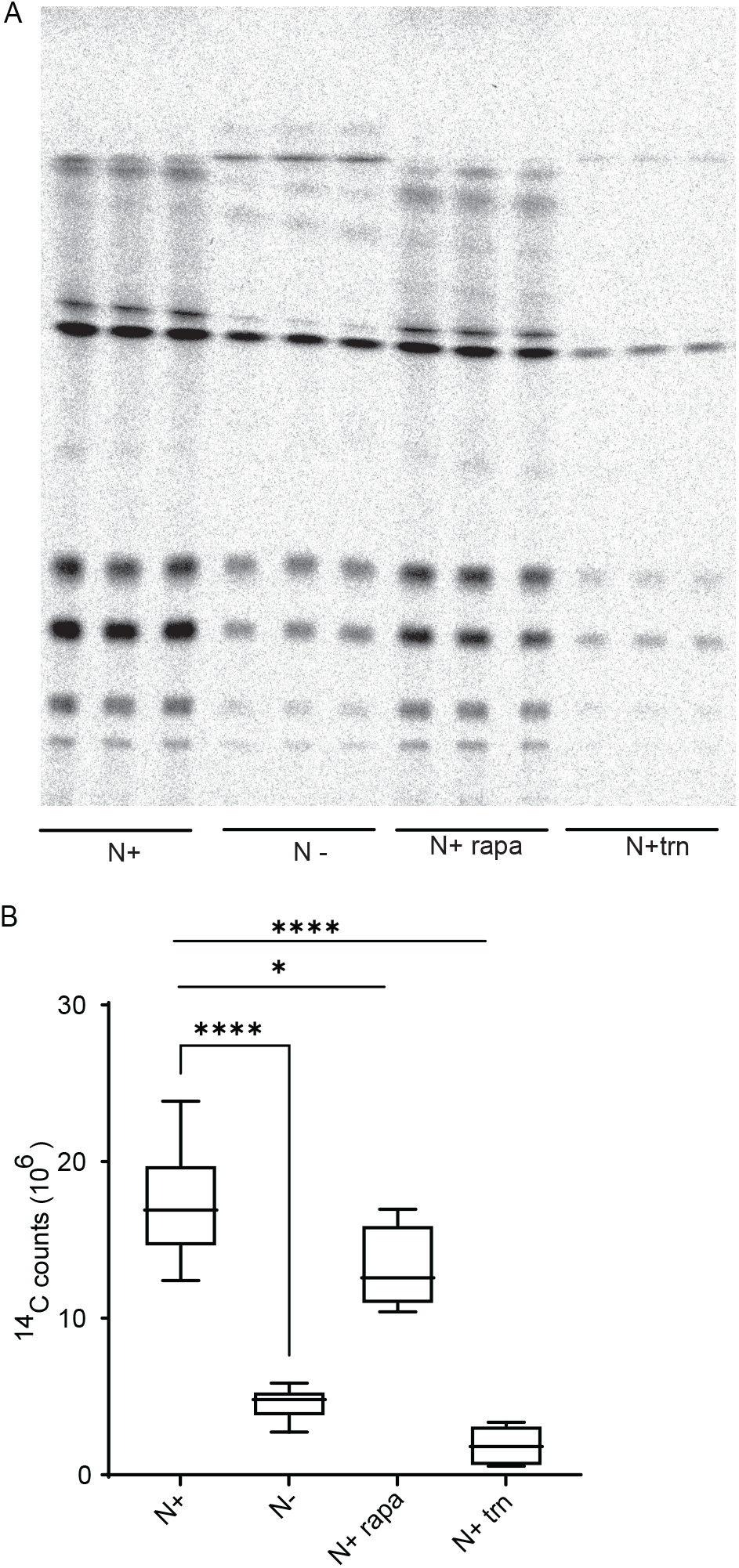
Regulation of fatty acid synthesis in *Chlamydomonas reinhardtii* under nitrogen limitation and TOR inhibition. A) TLC-autoradiograph of *C. reinhardtii* cells grown in N replete (N+), limited (N-), nitrogen replete with rapamycin (N+rapa) or torin1 (N+trn) fed with ^14^C acetate to track de novo synthesized fatty acid. B)^14^C-counts in fatty acid under nitrogen replete (N+), nitrogen deplete (N-), nitrogen replete with rapamycin (N+ rapa) and torin1 (N+ trn+). * indicate p-value confidence (student t-test).

## DISCUSSION

### Downregulation of fatty acid synthesis under N-limitation

Our study shows that the rate of fatty acid synthesis is decreased under N-limitation in the green algae *C. sorokiniana* and *C. reinhardtii*. This indicates that carbon flux into fatty acid synthesis is decreased during N-limitation in the green algal lineage. This information is crucial for any future biotechnological interventions to increase TAG accumulation in these organisms. This also raises the question, if inducing N-limitation-like conditions would be a means for channeling photosynthate to fatty acid, and in turn into oil production. Further, any strategy for enhancing oil production via the N-limitation response will have to account for the fact of decreased fatty acid synthesis under these conditions.

Nitrogen starvation mediated downregulation of fatty acid synthesis is conserved in the yeast *Saccharomyces cerevisiae.* In *S. cerevisiae,* under N-limitation, reduction in fatty acid synthesis is achieved by autophagy dependent specific degradation of FAS components (Shpilka et al., 2015). Prevention of FAS degradation decreased the viability under N-limitation. This indicates that *S. cerevisiae,* much like *C. sorokiniana* and *C. reinhardtii* actively decreases fatty acid synthesis under N-limitation. Conservation of decreased *de novo* fatty acid synthesis under N-limitation in evolutionarily diverse organisms, like microalgae and yeast, indicates that this is an essential adaptive response. When growth rate and cell division are limited by N-limitation, the demand for membrane glycerolipids will likewise be lower. When demand for membrane glycerolipids, a major sink for fatty acids, is lower, it would make sense for the cells to decrease the production of fatty acids. Further, fatty acid synthesis is an energy intensive process consuming NADPH and ATP molecules (Post-Beittenmiller et al., 1992). Decreasing carbon flux through this pathway would save energy (ATP) and reducing equivalents (NADPH). It is unclear if this is the only reason for the downregulation.

### TOR signaling under N-limitation

TOR signaling is an integral part of energy and nutrient sensing in eukaryotes. This is common to yeast, mammals, and plants (Shi et al., 2018). Multiple lines of evidence suggest the involvement of TOR signaling under N-limitation in microalgae. When treated with rapamycin under N-replete conditions, *C. reinhardtii* cells accumulate various amino acids within the first 5 minutes (Mubeen et al., 2018). This accumulation of amino acids is not from increased proteasomal activity but from active acquisition of nitrogen from the media via the activity of glutamine synthase (GS) and glutamine-2-oxoglutarate-aminotransferase. This active acquisition of nitrogen from the media is similar to an N-starvation response. Additionally, Mallen-Ponce et al., have shown that TOR activity is controlled by CO_2_ assimilation and this in turn controls the levels of certain amino acids in Chlamydomonas (Mallén-Ponce et al., 2022a). In a phosphoproteomic analysis of nitrogen starved cells of *C. reinhardtii*, Roustan et al. identified differential phosphorylation of ATG13, NNK1 and RSP6, known targets of TOR signaling (Roustan et al., 2017). This indicates that N-utilization/limitation and TOR signaling are linked in *C. reinhardtii*. Our work shows that under N-limitation in *C. sorokiniana*, there is a change in the abundance of individual proteins of the TOR complex (RAPTOR and TOR kinase) and differential phosphorylation of TOR kinase and RAPTOR. This coupled with significant overlap between the changes in the quantitative proteome and phosphoproteome under N-limitation and in TOR-inhibited cells indicates that TOR signaling is indeed affected under N-limitation. The relationship between TOR signaling and N-availability is not limited to microalgae. In Arabidopsis, nitrogen starvation leads to inactivation of TOR kinase and supplementation of select amino acids or inorganic nitrogen activates TOR kinase (Liu et al., 2021). In mammalian cells and *Saccharomyces cerevisiae,* TOR activity is controlled by amino acid availability via GTPases, RAB(A-D), and GRT1/2, respectively. Further, as mentioned earlier, in the fission yeast, *Schizosaccharomyces pombe,* deletion of TOR2 mimics nitrogen starvation (Matsuo et al., 2007).

It is not surprising that TOR signaling is controlled by N-availability in a variety of eukaryotes ranging from yeast and microalgae to land plants, since TOR functions as a central node in assimilating growth cues such as hormones and various nutrient availability (Xiong and Sheen, 2014), and nitrogen is an important macronutrient that regulates growth.

### TOR mediated regulation of fatty acid synthesis

In this study, we show that inhibition of TOR activity leads to a decrease in fatty acid synthesis thereby linking fatty acid synthesis to TOR signaling in the green plant (*Viridiplantae*) lineage. While there are transcriptional (Li et al., 2022) and protein level changes of ACCase and FAS complex members (this study), it is unclear if these are the only means by which TOR regulates fatty acid synthesis. It is also possible that TOR regulates the flux of carbon flow into the TCA cycle, in addition to the transcriptional and post-translational control of ACCase and FAS.

In mammalian systems, TORC1 regulates fatty acid synthesis at the transcriptional level by regulating SREBP1, a transcription factor that controls the expression and activities of ACCase and FAS complexes (Porstmann et al., 2008). It is interesting to note that TOR regulates fatty acid synthesis at the transcriptional level in evolutionarily distant organisms, such as mammals and microalgae. Evolutionary conservation of TOR mediated control of fatty acid synthesis can be attributed to two factors: TOR is a central regulator of metabolism that responds to many stress conditions such as nutrient limitation (Laplante and Sabatini, 2009), and fatty acid biosynthesis is a major sink for carbon and reducing equivalents. As mentioned earlier, decreasing fatty acid synthesis under nutrient limitation is in line with cellular needs under these stress conditions. TOR, being a central regulator of metabolism, coordinates the fatty acid synthesis process in response to external cues.

Knowledge of TOR functioning in the plant lineage has been lagging in comparison to that of mammalian and yeast cells, however in the past decade, the role of TOR kinase signaling as coordinator of metabolism in the green lineage has become clear (Robaglia et al., 2012; Caldana et al., 2013; Xiong and Sheen, 2014; Upadhyaya and Rao, 2019; O’Leary et al., 2020; Ingargiola et al., 2022; Mallén-Ponce et al., 2022b; Sharma et al., 2022). Our current work adds to the list of metabolic processes that are controlled by TOR signaling.

## CONCLUSION

In this work we have presented evidence to show that under N-limitation, microalgae like *C. sorokiniana* and *C. reinhardtii* reduce their rate of fatty acid synthesis, and that this downregulation is likely mediated by signal processing and transduction via the TOR kinase complex. This information will be useful in engineering microalgae to produce triacylglycerols and other storage compounds under nutrient replete growth conditions and may provide new insights into TOR-mediated control of fatty acid synthesis in land plants.

## MATERIALS AND METHODS

### Growth and N-limitation of C. sorokianiana and C. reinhardtii

*C. sorokianiana* and *C. reinhardtii* were grown in Tris Acetate Phosphate (TAP) media under continuous light with shaking (200rpm). *C. sorokiniana* was grown at 25°C while *C. reinhardtii* was grown at 22°C. TAP media was prepared as described by Gorman and Levine, 1965 (Gorman and Levine, 1965). In nitrogen deplete media (TAP-N), final concentration of ammonium chloride was 250 µM. For nitrogen starved cultures, parent cultures were grown for 2 days and washed thrice in appropriate media (TAP or TAP-N). The cultures were inoculated at a final concentration of 10^7^ cells/ml for *C. sorokiniana* and 5 x 10^6^ cells/ml for *C. reinhardtii*.

### Lipid extraction and total fatty acid assay

Lipid extraction was carried out using the Bligh and Dyer method (Bligh and Dyer, 1959). Glycerolipids were sequentially separated by Thin Layer Chromatography (TLC) on silica-G60 plates (EMD-Millipore) in a two-stage solvent system. First, the plate was developed for ∼2/3 of its length in chloroform, methanol, acetic acid, and water (85:12.5:12.5:3) to separate polar lipids. Next, the plate was dried and developed for the full length in petroleum ether, diethyl ether and acetic acid (80:20:1) to separate nonpolar lipids (triacylglycerol). For total fatty acid analysis, lipids extracted from 10^7^ cells were transesterified in methanolic HCl for 30 mins at 80°C to form Fatty Acid Methyl Esters (FAME). FAMEs were extracted into hexane by adding 1 ml hexane and 1 ml of 1 mM NaCl. The hexane phase was concentrated under a stream of nitrogen and quantified using GC-FID with Innowax column as previously described (Pflaster et al., 2014). 15:0 fatty acid was used as internal standard for quantification.

### 14C acetate labeling to measure de novo fatty acid synthesis

To measure *de novo* fatty acid synthesis, we carried out ^14^C-acetate feeding experiments. Cells were grown in appropriate growth conditions (N+ and N-) and then washed thrice in media lacking acetate (TP or TP-N) to remove any remaining acetate. Cells were then diluted to equal number of cells (10^7^ cell for *C. sorokiniana* and 5 x 10^6^ for *C. reinhardtii)* in 1 ml of TP / TP-N/ TP+ Torin 1/ TP+ rapamycin and 10^5^ cpm of ^14^C acetate (specific activity of 56mCi/mMol-supplied by ARC Inc.) was added and incubated for 45 mins. After 45 minutes the cells were washed thrice in appropriate media to remove any remaining ^14^C acetate. Lipid extraction and lipid separation was carried out as mentioned earlier. After drying, the TLC plates were exposed to a phosphoimager screen for 24 hours, and counts were read on a GE-Typhoon FLA 9500 scanner. Images were analyzed by ImageQuant TL v8.1. Total fatty acid level in each sample was calculated as a sum of all lipids in a given TLC lane.

### Inhibition of TOR signaling

In *C. sorokiniana* TOR activity was inhibited with 5 µM Torin 1 (EMD Millipore Corp.,). In *C. reinhardtii,* TOR kinase was inhibited using 1 µM rapamycin (Sigma Aldrich) or 5 µM Torin 1. To assay the effect of TOR inhibition on fatty acid synthesis, we incubated either *C. sorokiniana* or *C. reinhardtii* in TAP media with or without the inhibitors for 4 hours, and then labeled with ^14^C-acetate as described above.

### Protein preparation

For proteomic studies, samples were grown, treated, and collected as described (see results). Three replicates per condition were used for each treatment group. Protein samples for quantitative proteomic and phosphoproteomic analyses were prepared as follows: cells were ruptured by bead beating in lysis buffer (50 mM Tris-pH 8.0, 1 mM DTT, 0.5 mM PMSF and 2x PhosStop EASYpack-Roche) for 7 cycles with 30 seconds of beating at 400 rpm and 45 seconds on ice for each cycle. Glass beads of 0.5 mm diameter were used. Following the rupture, cells and debris were separated by spinning at 3000 rpm for 5 minutes at 4°C.

Proteins were precipitated and washed using the ProteoExtract™ Protein Precipitation Kit (MilliporeSigma, Burlington MA) and then redissolved in 8 M Urea, 0.2 M tris-HCl, pH 8.5 containing 2x PhosSTOP™ phosphatase inhibitor (Roche, Basel, Switzerland), 1x cOmplete™, EDTA-free Protease Inhibitor Cocktail (Roche) and 10mM DTT. Protein amounts were quantified using the CB-X™ protein assay (G-Biosciences, St Louis, MO). 1 mg of each sample was reduced at 37°C for 1 h and then alkylated with 20 mM iodoacetamide (20 min at RT in darkness), then quenched with an equimolar amount of DTT. Samples were diluted to 4 M urea with water and digested with 10 µg Lys-C (1:100 enzyme:substrate) at 37°C for 4 h. The urea was then diluted to 1 M by addition of 0.1 M tris-HCl, pH 8.5 and trypsin digestion carried out for 16 h at a ratio of 1:100. A further aliquot of trypsin (1:100) was added and digestion was carried out for a further 3 h. Digests were acidified with 20% TFA to pH 3, then desalted using 100 mg Sep-Pak® C18 reverse-phase SPE columns (Waters Corp, Milford, MA). Eluted samples were dried down and redissolved in 50 mM TEAB. A portion was set aside for analysis of the complete digest sample.

### Phosphoenrichment

For each sample, 0.75 mg of digested, desalted, dried peptide was dissolved in 2 M lactic acid, 60 % acetonitrile, 0.3 % TFA to 3 mg/mL and shaken vigorously with TiO_2_ beads (Titansphere, 5µm, GL Sciences, Tokyo, Japan) in a ratio of 1:5 sample: beads (w/w) for 20 h at 4°C. The beads were then washed with 3 x 100µL of the lactic acid binding solution and finally resuspended in 50µL and placed into 200 µL tips (Eppendorf, Hauppauge, NY) plugged with 2 layers of 3M™ C8 Empore™ membrane (ThermoFisher Scientific, Waltham, MA). For each sample, 100 µL of the same solution was spun through the beads at 3000 x g 3 times. The beads were then further washed with 3 x 200µL of 80 % acetonitrile, 0.4% TFA at 3000 x g. Bound phosphopeptides were then eluted 3 times with 100 µL ammonium hydroxide (5% v/v) into 1.5mL Lo-Bind tubes (Eppendorf). These were then frozen and lyophilized. A 2^nd^ elution of phosphopeptides was performed using 3 x 100 µL pyrrolidine (5% v/v) at 1000 x g and the samples were frozen and lyophilized. Both eluates for each sample were pooled and desalted using an Oasis µElution plate (Waters Corp), dried down and then redissolved for analysis by LC-MS/MS.

### LC-MS/MS analysis of the proteome and phosphoproteome

Peptide and phosphopeptide samples were analyzed by LC-MS/MS on an RSLCnano system (ThermoFisher Scientific) coupled to a Q-Exactive HF mass spectrometer (ThermoFisher Scientific). The samples were first injected onto a trap (Acclaim PepMap™ 100, 75µm x 2 cm, ThermoFisher Scientific) for 2.8 min at a flow rate of 5 µL/min, 1.5% acetonitrile, 0.2% formic acid before switching in-line with the main column. Separation was performed on a C18 nano column (Acquity UPLC® M-class, Peptide CSH™ 130A, 1.7µm 75µm x 250mm, Waters Corp) at 260 nL/min with a linear gradient from 5-32% over 96 min. The LC aqueous mobile phase contained 0.1% (v/v) formic acid in water and the organic mobile phase contained 0.1% (v/v) formic acid in 80% (v/v) acetonitrile. Mass spectra for the eluted peptides were acquired on a Q Exactive HF mass spectrometer in data-dependent mode using a mass range of m/z 375–1500, resolution 120,000 for the MS1 peptide measurements. Data-dependent MS2 spectra were acquired by HCD fragmentation with a normalized collision energy (NCE) set at 28%, AGC target set to 1 x 10^5^, 15,000 (for phosphopeptides at 30,000) resolution, intensity threshold 1 x 10^5^ and a maximum injection time of 118 ms (86 ms for phosphopeptides). Dynamic exclusion was set at 45 sec for the complex digest and 30 sec for the phosphopeptide analysis to help capture phospho isomers.

### Data analysis

Data were analyzed in Proteome Discoverer 2.2 software (ThermoFisher Scientific) and searched using Mascot 2.6.2. The databases searched were the common contaminants database cRAP (116 entries, www.theGPM.org) and in-house *Chlorella sorokiniana* gene catalog with13,906 entries. Gene identifiers and sequence data used for the analysis are available at NCBI under BioProject PRJNA34632. Methionine oxidation, protein N-terminal acetylation, deamidation of glutamine, carbamidomethylation of cysteine and Ser/Thr and Tyr phosphorylation were set as variable modifications. A maximum of two trypsin missed cleavages were permitted and the precursor and fragment mass tolerances were set to 10 ppm and 0.02 Da, respectively. Peptides were validated by Percolator with a 0.01 posterior error probability (PEP) threshold. The data were searched using a decoy database to set the false discovery rate to 1% (high confidence). The localization probabilities of the PTMs were obtained using ptmRS (Taus et al 2011). The peptides were quantified using the precursor abundance based on intensity. The peak abundance was normalized using total peptide amount. The peptide group abundances are summed for each sample and the maximum sum for all files is determined. The normalization factor used is the factor of the sum of the sample and the maximum sum in all files. The protein ratios are calculated using a pairwise ratio-based method. The protein ratios are calculated as median of all possible pairwise ratios between the replicates of all connected peptides. To compensate for missing values in some of the replicates, the replicate-based resampling imputation mode was selected. The significance of differential expression is tested using an ANOVA test which provides adjusted p-values using the Benjamini-Hochberg method for all the calculated ratios.

### Multiple Sequence Alignment

TOR-kinase sequence for different algae were collected from Phycocosm (Grigoriev et al., 2021) and Multiple sequence alignment was performed using Clustal Omega (Madeira et al., 2022).

### Statistical analysis

All statistical analysis for biochemical and growth assays were performed using Graphpad prism v9.

## Supporting information

Supplement_proteome_CSr

## Acknowledgement

This work was supported by National Science Foundation award number EPS-1004094 (to WRR.), NASA award number 80NSSC17K0737 (to WRR) and startup funding from the University of Nebraska-Lincoln College of Arts and Sciences and School of Biological Sciences (to WRR). The Proteomics & Metabolomics Facility (RRID:SCR_021314) and instrumentation are supported by the Nebraska Research Initiative.

## Author Contributions

JV conceived the hypothesis, designed, and performed the experiments and wrote the manuscript. MN and SA performed the proteomic, phosphoproteomic analysis and edited the manuscript. AM performed the fatty acid analysis. NW and WRR provided critical insights and edited the manuscript.

---

**Fig. S1.**
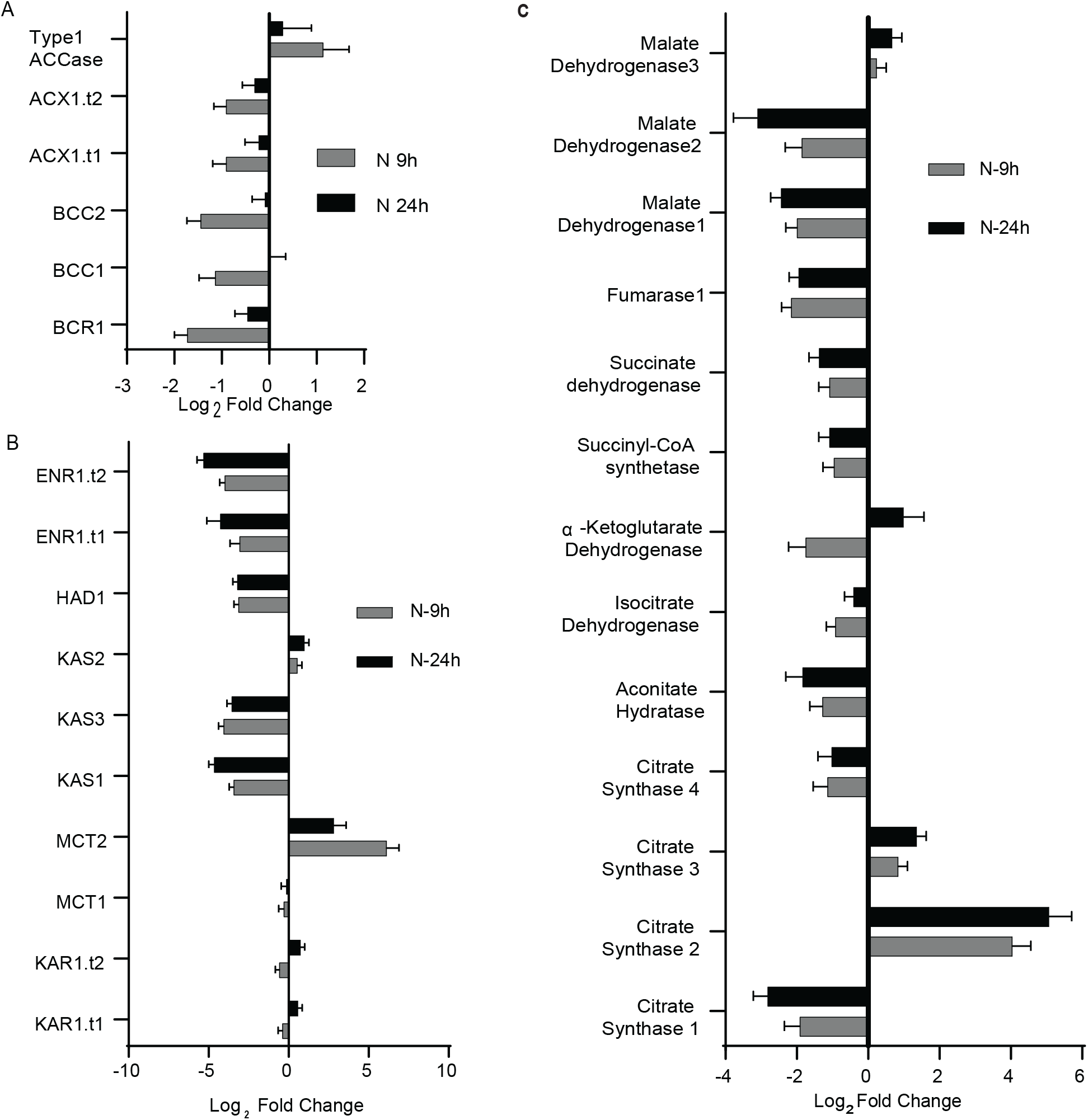
A) Expression profile of ACCase complex genes under nitrogen limitation in C. sorokiniana at 9 and 24 hours after inoculation. BCR1: Biotin carboxylase, BCC: Acetyl-CoA biotin carboxyl carrier protein, ACX: Carboxyltransferase, type1 ACCase: homomeric Acetyl CoA Carboxylase. B) Expression profile of FAS complex genes under nitrogen limitation in C. sorokiniana. KAR: 3-Oxoacy-ACP reductase, MCT: Malonyl-CoA: ACP transacylase, KAS: 3-Ketoacyl-ACP-synthase, HAD: 3-Hydroxyacyl-ACP dehydratase, ENR: Enoyl-[acyl carrier protein] reductase. C) Expression of TCA cycle genes under nitrogen limitation. Fold change is relative to cells grown under nitrogen replete media at the appropriate time point.

**Fig S2.**
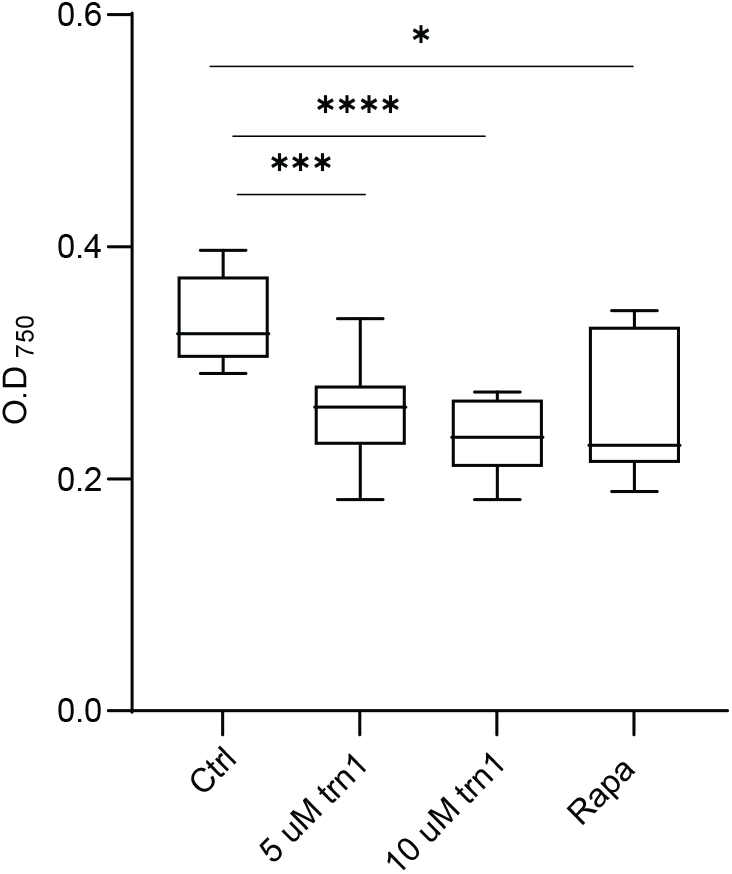
Inhibition of growth of *C. sorokiniana* by Torin 1 or rapamycin. *C. sorokiniana* cultures were inhibited with either 5,10 uM Torin 1 (trn1) or 10 uM rapamycin (Rapa). The graph indicates O.D after 1 day of growth. * indicate statistical significance of a Welch’s t test (n=9).

**Figure S3.**
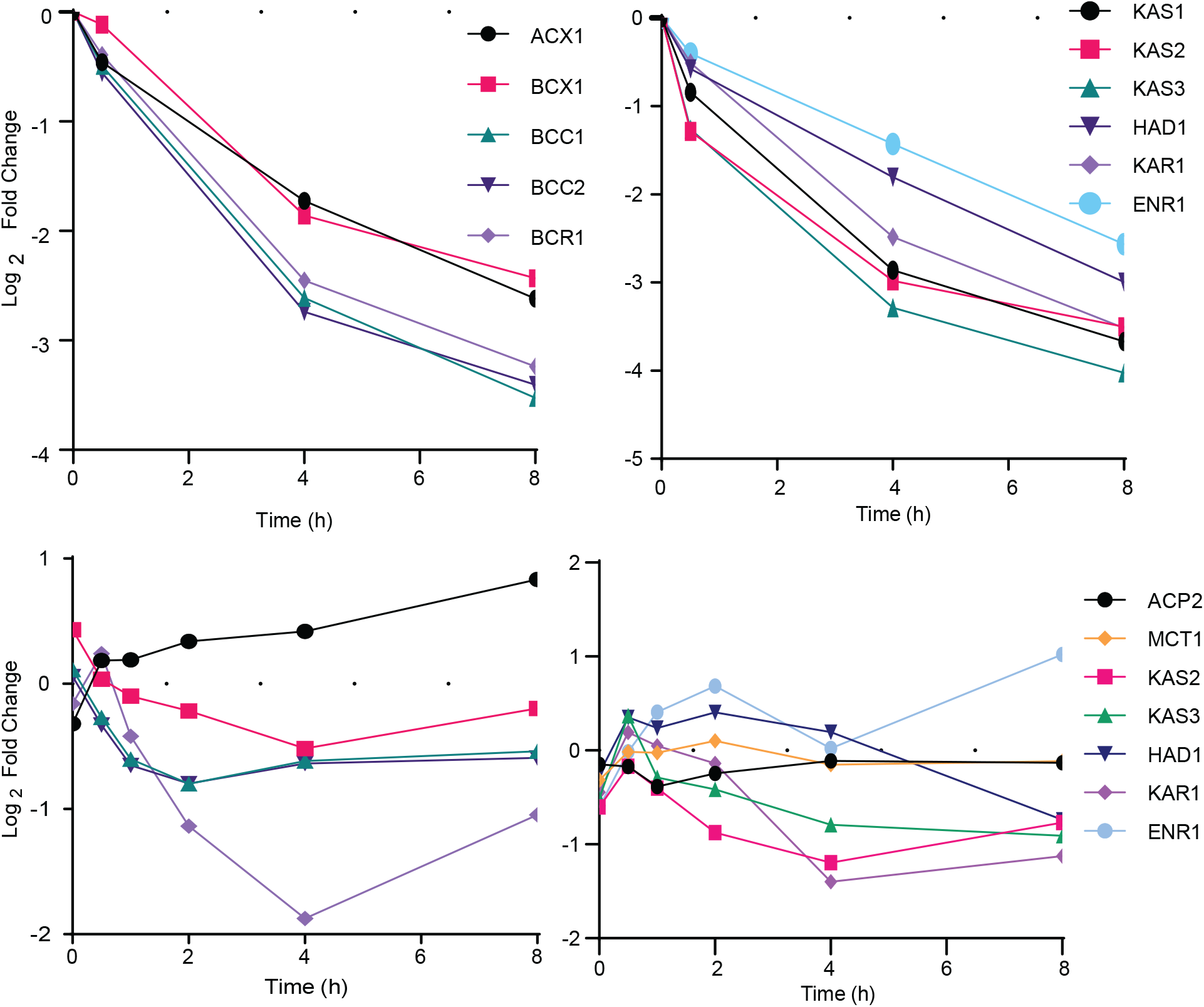
TOR regulation of fatty acid synthesis in *Chlamydomonas reinharditii*. A) Expression of ACCase complex genes under nitrogen limitation. B) Expression of FAS complex genes under nitrogen limitation. Data for this analysis was from https://phytozome.jgi.doe.gov/pz/portal.html. C) Expression of ACCase complex genes when treated with rapamycin. D) Expression of FAS complex genes when treated with rapamycin. Data for this analysis was from Kleessen et al., 2015.

